# Inhibition of interferon signalling improves rabbit calicivirus replication in biliary organoid cultures

**DOI:** 10.1101/2025.03.28.645886

**Authors:** Elena Smertina, Megan Pavy, Nias Y. G. Peng, Omid Fahri, Maria Jenckel, Tanja Strive, Michael Frese, Ina L. Smith

## Abstract

The *Rabbit haemorrhagic disease virus* (RHDV) was discovered 40 years ago. This highly pathogenic virus threatens the integrity of ecosystems in the European rabbits’ native range, while in Australia, it is used as a biocontrol tool to manage overabundant populations of feral rabbits (*Oryctolagus cuniculus*). Little is known about the life cycle of this virus due to the absence of a reliable cell culture system. In 2023, we developed a rabbit liver-derived organoid cell culture system that supports RHDV replication but is unable to sustain serial passaging in culture. Here, we report that the interferon signalling pathway inhibitor Ruxolitinib increases virus replication in organoid-derived monolayer cells and, for the first time, enables the serial passaging of RHDV in cell culture. Four consecutive passages were achieved with viral titres reaching the concentration of the initial virus stock as measured by RT-qPCR. Immunofluorescence analysis showed that more cells are infected in the presence of Ruxolitinib. Furthermore, we noted that cells grew faster and formed healthier monolayers in the presence of the interferon inhibitor. To determine the cellular composition of the monolayers, we used single-cell RNA-sequencing, revealing that our organoids consist largely of RHDV-permissive cholangiocytes.

**IMPORTANCE:** In this work, we describe the use of interferon inhibitor to enhance the permissiveness of our recently developed rabbit liver-derived organoid cell culture to RHDV infection. Interferon inhibitors offer a simple and cost-effective approach to increase the replication of difficult-to-grow viruses in culture systems that are not interferon-deficient.

## INTRODUCTION

Rabbit haemorrhagic disease viruses (RHDVs) are highly pathogenic, positive sense, single-stranded RNA virus in the genus *Lagovirus*, family *Caliciviridae*, that infect lagomorphs. There are two major genotypes; RHDV1 infects rabbits while RHDV2 also infects other lagomorphs, such as hares. Both genotypes are hepatotropic, and an infection usually leads to peracute liver disease, characterized by necrosis, haemorrhages, and death within 72 h. The highly virulent nature makes RHDV an efficient biocontrol agent for the introduced European rabbit population in Australia and New Zealand (reviewed in (1)).

Although RHDV1 was discovered in 1984 (2)and has been used in Australia as a biocontrol agent for rabbits since the mid-1990s, knowledge of many aspects of the viral life cycle is limited. For example, the virus entry receptor is unknown, and the functions of several of its non-structural proteins have not yet been determined. This can mainly be attributed to the inability to efficiently grow rabbit caliciviruses in cell culture, a problem that is frequently encountered with hepatotropic viruses, such as the human hepatitis B and C viruses (reviewed in (3)). Hepatocytes, the main target cells for these viruses, are highly differentiated cells that are difficult to grow and maintain in cell culture. Recently, we established rabbit liver-derived organoid cultures that support the replication of both RHDV1 and RHDV2 (4). However, only a small proportion of organoid cells produced detectable amounts of viral antigen and attempts to passage the virus in liver organoids were not successful. Thus, further improvements were needed to generate a more robust cell culture system.

Virus infections in cell culture often trigger interferon (IFN) responses mediated by IFN type I (IFN-α, IFN-β and others) and IFN type III (IFN-λ). IFN type II (IFN-γ) is usually not a factor that limits virus replication in cell culture as it is predominantly produced by immune cells (reviewed in (5)). In immune competent cells, the production of IFN and the subsequent expression of IFN-induced genes can inhibit or completely block virus multiplication. For example, human noroviruses (family *Caliciviridae*) induce innate immune responses in human intestinal organoids via IFN types I and III signalling (6, 7). As IFNs regulate gene expression through the Janus kinase (JAK) signal transducer and activator of transcription (STAT) signalling pathway (reviewed in (8)), it is not surprising that genetically engineered STAT1-knockout intestinal organoids demonstrate an increased permissiveness to human norovirus infection (7).

Ruxolitinib (Rux), a small-molecule JAK1 and JAK2 inhibitor, inhibits early stages of all types of IFN signalling pathways by competing with adenosine triphosphate (ATP) for the ATP-binding pocket of these kinases (9). Rux and similar IFN inhibitors increase the replication of a range of viruses when used as a supplement in cell culture media (6, 10–12). Apart from inhibiting IFN-mediated responses, Rux can induce apoptosis in cancer cells (13, 14), which makes it difficult to reliably predict the benefit of its use in cell culture.

In this work, we aimed to optimize the rabbit liver-derived organoid system to enable passaging of lagoviruses and the production of high-titered virus stocks, by assessing the ability of the IFN signalling inhibitor Rux to increase RHDV replication. Furthermore, using single-cell RNA-sequencing, we determined the cellular composition of the organoid-derived monolayers, which revealed that our organoids contain predominantly cholangiocytes.

## MATERIALS AND METHODS

### Establishment and culturing of rabbit liver organoids and organoid-derived monolayers

The rabbit organoid (spheroid) cultures and organoid-derived monolayers were established and passaged as described previously (4). Modifications included the use of Rux (Selleckchem or Sapphire Bioscience) in cell culture media (with a concentration of 4 μM if not otherwise specified) starting from the first seeding of isolated stem cells unless otherwise specified. Rux stock solution was prepared by reconstitution in dimethyl sulfoxide (DMSO) to a 10 mM stock concentration.

### Viruses

RHDV1 (genotype GI.1cP-GI.1c according to Le Pendu *et al*. (15); GenBank accession number KT344772) and RHDV2 (genotype GI.1bP-GI.2; GenBank accession number MW467791) stocks were prepared by the Elizabeth Macarthur Agricultural Institute (Menangle, NSW, Australia) from semi-purified liver homogenates after amplification in domestic rabbits, and the rabbit infectious dose (ID_50_) was titrated in rabbits as described previously (16, 17). Freeze-dried virus stocks were reconstituted in unsupplemented advanced DMEM/f12 (Sigma-Aldrich) and stored at −80°C.

### Inoculation of biliary organoid monolayer cultures

The organoid-derived monolayers were seeded on Geltrex-(Gibco) or Matrigel-(Corning) coated surfaces. The coating was performed using a 2% solution of Geltrex or Matrigel in Dulbecco’s phosphate-buffered saline (DPBS; Sigma-Aldrich) for 2 h at 37°C. The seeding density had been determined empirically and is 80,000 cells per well for an 8-well chamber slide (Thermo Scientific). The cells were inoculated in triplicates with either RHDV1 or RHDV2 at a MOI of 0.01. The inoculum was diluted in DMEM high glucose medium (Sigma-Aldrich) supplemented with 2% heat-inactivated FCS (Thermo Scientific). Mock-infected cells and cells incubated with heat-inactivated RHDV1 and RHDV2 (65°C for 15 min) were used as controls. After 1 h at 37°C in a 5% CO_2_ incubator, inoculum was removed, and cells were washed with DPBS three times. Fresh culture medium (composition as described in (4)) was then added, and the cells were incubated for 1, 48 or 72 h, as specified.

### RNA extraction and RT-qPCR

Total RNA was extracted using the NucleoSpin RNA Mini kit (Macherey-Nagel) or KingFisher Flex instrument (Thermo Scientific) according to the manufacturer’s instructions, and viral RNA was quantified using a universal RT-qPCR assay targeting a conserved region of the lagovirus VP60 coding sequence as described previously (Hall et al., 2018) using the SensiFAST SYBR No-ROX kit (Bioline Reagents, Meridian Bioscience) on a CFX96 Touch real-time PCR instrument (Bio-Rad). A standard curve was utilized for absolute quantification; ‘no template’, previously quantified positive control and extraction controls were included in each analysis plate. Each sample was processed using three technical replicates. The data was analyzed with CFX Maestro Bio-Rad software, and results reported as log_10_ capsid gene copies per well.

To analyze IFN-induced gene expression, primers were designed to cover intron-spanning regions using Primer Express 2.0 (Thermo Scientific) or PrimerQuest Tool (Integrated DNA Technologies) software (Supplementary Table 1). RT-qPCRs were performed using the SensiFAST Probe No-ROX kit (Bioline Reagents, Meridian Bioscience) on a CFX96 Touch real-time PCR instrument (Bio-Rad) as follows: reverse transcription at 45°C for 10 min, followed by initial denaturation at 95°C for 2 min, then 40 cycles of 95°C for 5 s and 60°C for 20 s. A ‘No template’ control was added with each run. Expression of the gene of interest was normalized to expression levels of the housekeeping gene glyceraldehyde-3-phosphate dehydrogenase (*GAPDH*). Fold change was calculated using ΔC_T_ method (18).

### Immunofluorescence staining and analysis

Monolayer cultures grown in chamber slides were fixed with 100% ice-cold acetone at −20°C for 12 min, washed twice with DPBS, permeabilized with 0.25% Triton X-100 (Sigma-Aldrich) in DPBS for 10 min at room temperature, washed with DPBS three times, and blocked with DPBS containing 5% BSA and 0.1% Tween-20 (DPBST) for 60 min. For virus capsid staining, the lagovirus-specific anti-VP60 mouse monoclonal antibody 13C10 (25 µg/ml; Monoclonal Antibody Facility of the Institute for Medical Research, Perth, Australia) (19) was used at a 1:100 dilution in 1% BSA in DPBST. The slides were then incubated overnight at 4°C. The cells were then washed three times with DPBS, and secondary antibodies conjugated to Alexa Fluor 488 or 555 (Abcam; ab150113 and ab150114, respectively) were added (diluted at 1:300 in 1% BSA in DPBST). The slides were incubated at room temperature for 60 min in the dark, washed three times with DPBS, and stained with DAPI (1 mg/ml; Sigma-Aldrich) diluted at 1:500 in sterile water. The cover glass was mounted with FluoroShield (Sigma-Aldrich). Fluorescence images were obtained using a Zeiss AxioImager M2 microscope equipped with plan apochromat 20x NA=0.8 and 10x NA=0.45 objectives, Colibri 7 fluorescence LED illumination and Zeiss Axiocam 712 colour CCD camera (Carl Zeiss). Zeiss filter sets 96 HE (excitation BP390/40; beam splitter FT 420; emission BP450/40) with LED excitation band 370–400 nm, and 38 HE (excitation BP470/20; beam splitter FT495; emission BP 525/25) with LED excitation band 450–488 nm were used to image DAPI and VP60 fluorescence respectively. Image capture was carried out using ZEN blue v3.2 (Carl Zeiss). Quantitative image analysis was completed using FIJI/ImageJ v. 1.54 (20). Briefly, fluorescence images were converted to 8-bit grayscale, an identical threshold applied, and the FIJI “Analyze particles” function was used to quantify the total number of cells and VP60 expressing cells.

### Data visualization and statistical analyses

The quantitative data were processed and visualized in R (R core team, 2024) using packages ggplot2 (v3.5.1) (21), patchwork (v1.3.0) (22), ggpattern (v1.1.4) (23), dplyr (v1.1.4) (24), stringr (v1.5.1) (24), rstatix (v0.7.2) (25), magrittr (v2.0.3) (26), and svglite (v2.1.3) (27). Statistical differences were assessed using one-way or two-way ANOVA (as specified) followed by Tukey’s honestly significant difference (HSD) test.

### Single-cell RNA-sequencing

Rabbit liver organoid-derived monolayer cells were seeded on Geltrex-precoated 6-well plates using media supplemented with Rux and incubated overnight. The next day, the cells were detached using TrypLE (Sigma-Aldrich), counted using a hemocytometer, and passed through a 40-µm cell strainer. The cells were resuspended in 2% FBS in DPBS at 1 million/ml. Prior to FACS sorting, propidium iodide (50 μg/ml, BD Biosciences) was added (0.1 μg/million cells) to discriminate between live and dead cells. Hoechst (BD Biosciences) staining was used at 4 μg/ml for sorting single cells and excluding duplets and clumps. The cells were then incubated for 15 min at 37℃ prior to sorting with the FACSAria Fusion (BD Biosciences) instrument (John Curtin School of Medical Research, Australian National University (ANU)). A total of 84,000 live single cells were collected and processed using 10x Chromium (10x Genomics) single-cell sequencing library preparation following the manufacturer’s protocol (Biomolecular Resource Facility, ANU). The resulting cDNA was sequenced with Illumina NextSeq 2000 (P2 100 cycle flow cell) instrument (Biomolecular Resource Facility, ANU). The raw reads were processed using Cell Ranger software, version 7.1.0 (28). The reads were mapped to *Oryctolagus cuniculus* genome (NCBI RefSeq GCF_009806435.1). The marker gene expression analysis and visualisation were performed with Python package scanpy, version 1.10.4 (29).

## RESULTS

### Rabbit liver organoid-derived monolayers are predominantly of biliary origin

Previous analyses using RT-qPCR and immunofluorescence staining revealed that the majority of the organoid-derived monolayer cells are not hepatocytes (4). Although our earlier work suggested that the cells were of biliary origin, a precise analysis was challenging due to the lack of rabbit cell marker-specific antibodies. Here, to identify the cellular composition of the organoid-derived monolayer cells with greater precision, we used single-cell RNA-sequencing (Fig. 1).

**FIG 1.**
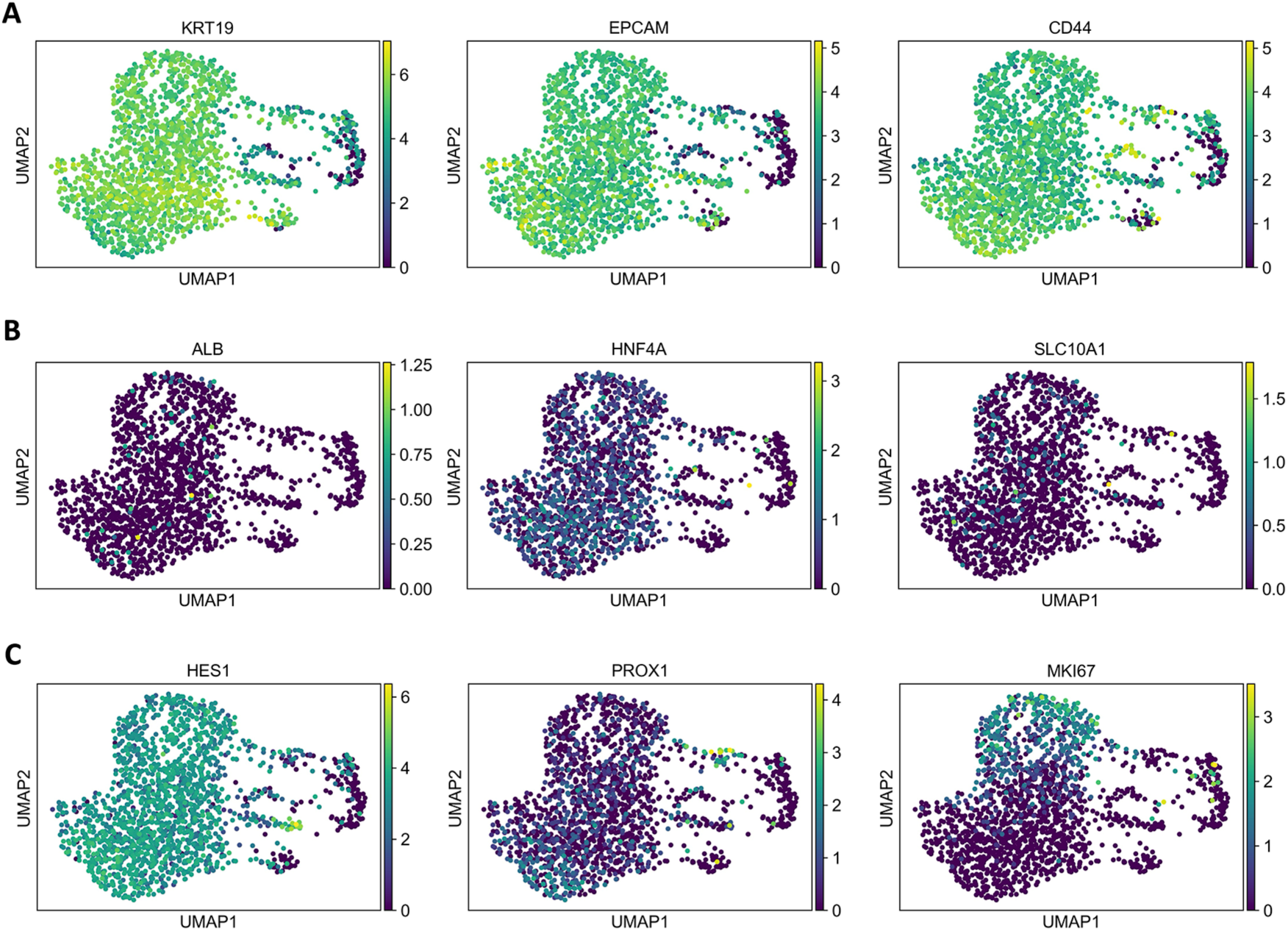
Uniform manifold approximation and projection (UMAP) plots of normalized marker gene expression in rabbit liver monolayers. Each dot represents a cell, and the color indicates the level of gene expression, with lighter colors indicating higher level of expression. (**A**) Expression of the cholangiocyte-specific markers cytokeratin-19 (KRT19), epithelial cell adhesion molecule (EPCAM), and epithelial cell marker cluster of differentiation 4 (CD44). (**B**) Expression of the hepatocyte-specific markers albumin (ALB), hepatocyte nuclear factor 4 alpha (HNF4A), and sodium-taurocholate co-transporting polypeptide (SLC10A1). (**C**) Expression of the early development transcription factor hairy and enhancer of split-1 (HES1), Prospero homeobox protein 1 (PROX1), and marker of proliferation Kiel 67 (MKI67).

We observed increased levels of expression of cholangiocyte-specific genes compared to hepatocyte markers, in line with our previous results (4). For example, EPCAM and KRT19 were highly expressed (Fig. 1A), a characteristic feature of cholangiocytes (30). High expression of epithelial progenitor cell marker CD44 was also observed, suggesting that the analyzed cells are immature cholangiocytes. Mature hepatocyte markers (ALB, HNF4A, and SLC10A1) were expressed at low levels (Fig. 1B), confirming that the monolayer cells are predominantly immature bile duct cells. The conclusion was further corroborated by the high expression of the early development transcription factor HES1, which is essential for tubular bile duct formation (31), and low expression levels of PROX1, responsible for hepatocyte proliferation and migration (32).

We observed a mostly uniform expression of cell-type specific markers in our monolayer cultures with the exception of the proliferation marker (MKI67) (Fig. 1C). Only a subset of cells expressed high levels of MKI67, which can be a consequence of different cell cycle stages.

### Rux enhances RHDV2 replication in rabbit liver monolayers

The small-molecule IFN inhibitor Rux was tested for its ability to boost RHDV2 replication. Rux is a pan-IFN inhibitor that affects the phosphorylation of STAT1, a key component of the IFN-induced signalling cascade. In a first experiment, organoid-derived monolayers were treated with Rux for 48 h prior to infection, and the inhibitor was tested at two concentrations, 1 and 4 μM. Cells grown in Rux-free media or media supplemented with the equivalent volume of DMSO were used as controls. Both the cells and supernatant were collected at 1 and 72 h post infection (hpi) and virus replication was quantified using RT-qPCR. Similar (∼1 log_10_) increases in RHDV2 replication were observed in cells cultured with 1 and 4 μM Rux compared to untreated or DMSO-supplemented cells at 72 hpi (Fig. 2A).

**FIG 2.**
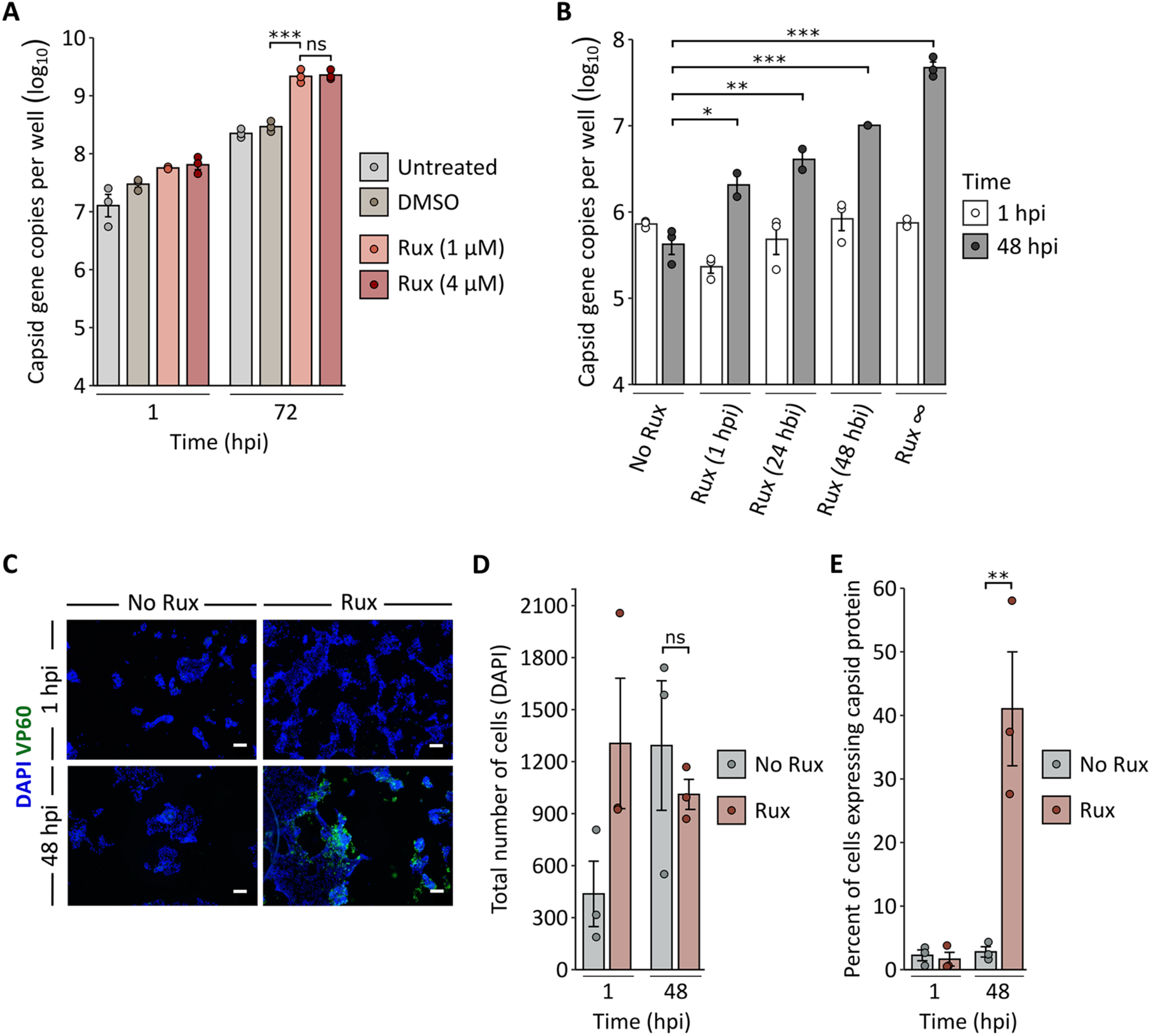
Effect of Rux on RHDV2 replication in rabbit liver organoid-derived monolayers. (**A**) Rabbit liver monolayers supplemented with Rux 48 h prior to infection with RHDV2 at a MOI of 0.01. Virus replication was quantified using RT-qPCR at 1 and 72 hpi. (**B**) Rux was added to the media either before infection with RHDV2 (∞, 48, 24 hbi), or after infection (1 hpi). Virus replication was quantified using RT-qPCR at 1 and 48 hpi. Individual data points represent three biological replicates averaged from three technical replicates for each; columns and error bars represent the mean values and the standard error of the mean, respectively (A and B). (**C**) Immunofluorescence staining of RHDV2-infected rabbit liver monolayer cells in the presence or absence of Rux. Monolayer cells grown in chamber slides were stained with antibodies against the lagovirus capsid protein (VP60, green). The nuclei were counterstained with DAPI (blue). Scale bar, 100 µm. (**D**) Cells were counted based on DAPI fluorescence. (**E**) VP60 expression was quantified as a ratio of VP60-positive cells to the total number of cells. Individual data points represent results of the analysis for one image. Columns and error bars represent the mean values and the standard error of the mean, respectively (D and E). Significance is denoted by asterisks (\**p*<0.05, \*\**p*<0.01, \*\*\**p*<0.001) and was calculated using one-way (A and B) or two-way (D and E) ANOVA followed by Tukey’s HSD test.

To characterise the effect of Rux over time and determine the optimal supplementation time, we investigated whether culturing of cells in the presence Rux ahead of the infection would boost virus growth further compared to shorter incubation periods. To this end, 4 μM Rux was added to the culture media either from the first cell passage of freshly cultivated rabbit liver cells, 48 and 24 h prior to infection, or 1 h after RHDV2 inoculation. For each group, mock-infected cells that received the corresponding Rux treatment but not infected with RHDV2 were included as controls (not shown). A significant increase in virus replication was detected in all treatment groups compared to cells that did not receive Rux (Fig. 2B). Furthermore, we noted that a prolonged culturing of cells in the presence of Rux tended to result in higher virus replication (Fig. 2B).

Next, we analyzed the effect of Rux at a cellular level by immunofluorescence staining of RHDV2-infected cells cultured with or without 4 μM Rux. A significant increase in the number of RHDV2-infected cells cultured with Rux was detected at 48 hpi (Fig. 2C, E). Although an equal cell number was seeded in each well, more cells were observed in Rux-supplemented samples at 1 hpi compared to Rux-free wells (Fig. 2C, D), suggesting that incubation with Rux promotes the growth of monolayer cells. Of note, cell numbers at 48 hpi were similar (Fig. 2D), indicating that the difference in number of VP60 (capsid) expressing cells cannot be attributed to the difference in the total cell number at that timepoint (Fig. 2E). Rather, the observed difference is due to a reduced permissiveness and/or a lower number of cells at 1 hpi in Rux-free samples.

### Rux inhibits the expression of IFN-induced genes in infected rabbit liver monolayers

To investigate the effect of Rux on IFN signalling in RHDV-infected cells, we quantified the expression of three IFN-induced genes, i.e., guanylate-binding protein 1 (*GBP1*), IFN-induced protein with tetratricopeptide repeats 1 (*IFIT1*), and myxovirus resistance protein 1 (*MX1*). The expression of *GBP1* and *IFIT1* was significantly higher in cells infected with RHDV2 in the absence of Rux, than in Rux-free cells (Fig. 3). For *MX1*, the difference was not statistically significant (Fig. 3). Of note, the expression of *GBP1*, *IFIT1*, and *MX1* in control samples (mock-infected and infected for 1 h) was generally below the expression of the housekeeping gene *GAPDH* that was used for normalization. Our findings indicate that RHDV infection triggers the IFN response, and that there is no constitutive expression of IFN-induced genes in uninfected cells.

**FIG 3.**
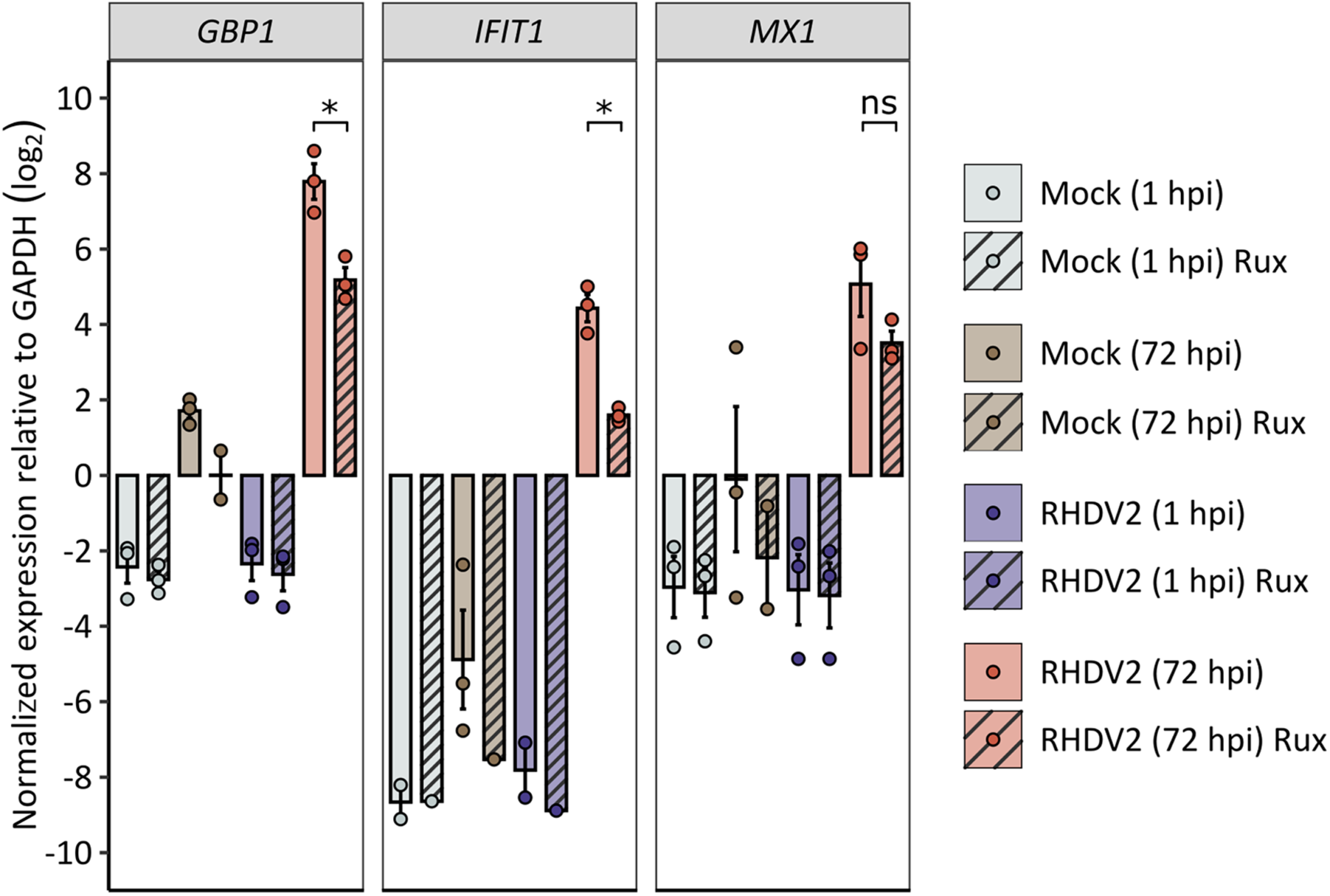
Quantification of IFN-induced gene expression in rabbit liver monolayers. Expression of *GBP1*, *IFIT1*, and *MX1* were measured using RT-qPCR in mock-infected (grey and brown) and RHDV2-infected (purple and red) monolayer cells at 1 and 72 h post infection (hpi). Striping indicates the addition of Rux 48 h prior to infection. Gene expression is presented as log_2_ fold change (ΔCT) relative to the expression of a housekeeping gene (*GAPDH*). Means were calculated from three independent biological replicates with three technical RT-qPCR replicates each. Error bars represent the standard error of the mean. Asterisks indicate statistical significance (ns, not significant; \**p*<0.05), as determined by one-way ANOVA followed by Tukey’s HSD test.

### Rux enables serial RHDV passaging in rabbit liver monolayers

As attempts to passage RHDV in rabbit liver monolayers in the absence of IFN inhibitors were not successful (Supplementary Fig. 2), we tried passaging the virus in the presence of Rux. To that end, monolayer cells were inoculated with 0.01 MOI of either RHDV1, RHDV2, heat-inactivated viruses or mock-infected, and incubated for 72 h. Aliquots of supernatants were diluted 1:10 with fresh media and used as inoculum for the next passage; we performed a total of 4 passages and quantified the virus genome copy number for each passage at 1 and 72 hpi. In every passage, the titre in the supernatant almost reached the initial titre of the inoculum demonstrating that the interferon inhibitor Rux allowed a productive passaging of both RHDV1 and RHDV2 (Fig. 4).

**FIG 4.**
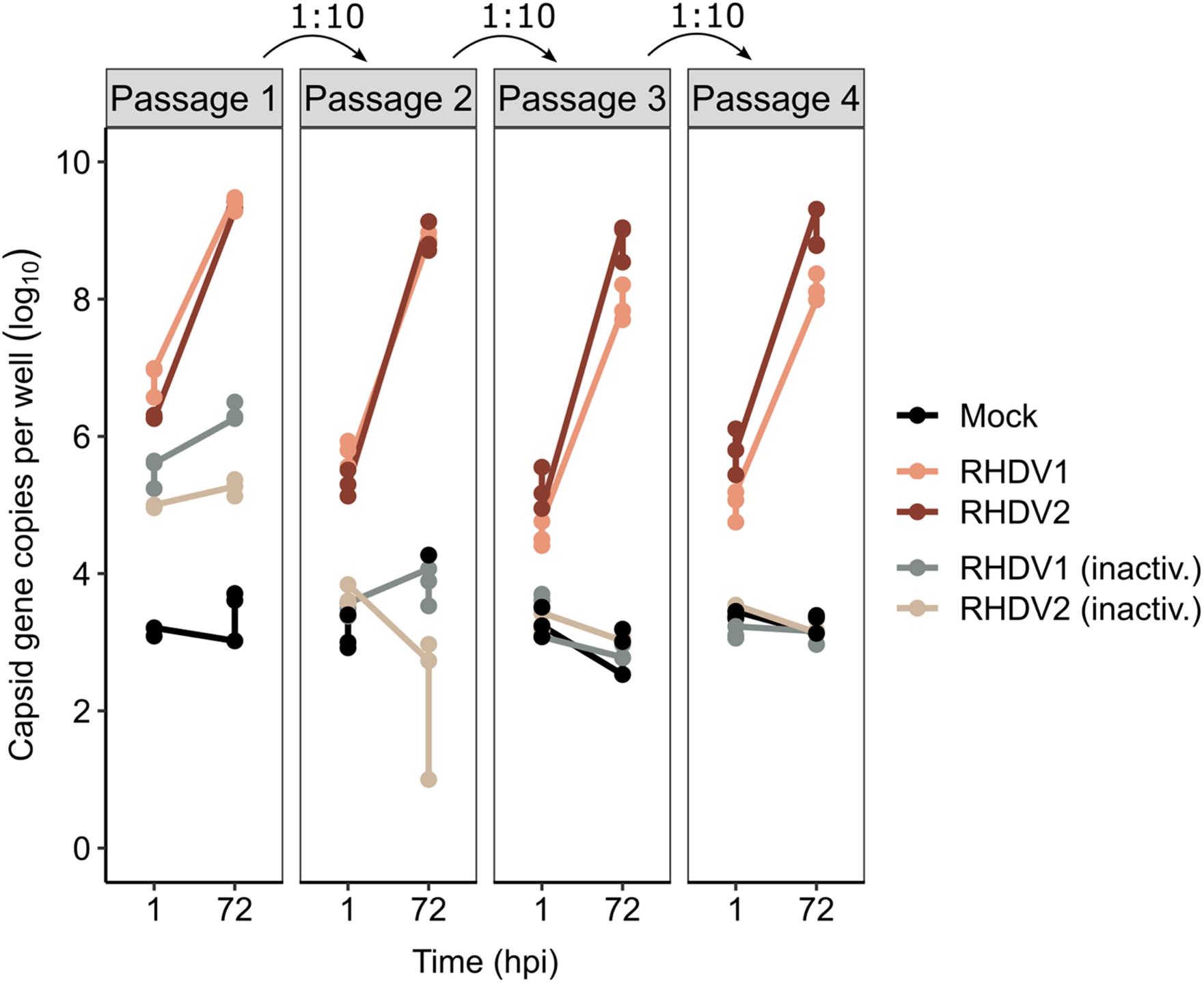
Passaging of RHDV in rabbit liver monolayers in the presence of Rux. The cells were either mock-infected (Mock) or infected with RHDV1, RHDV2, or heat-inactivated viruses (inactiv.). The capsid gene copy numbers were quantified at 1 and 72 h post infection (hpi). The supernatant from passage one diluted at 1:10 was used as inoculum for passage two and so on until passage four. Individual data points represent three biological replicates averaged from three technical replicates for each.

## DISCUSSION

The lack of a reliable culture system has long hampered research on rabbit caliciviruses. Our previously reported organoid culture system allows the growth of RHDV in cell culture, but virus infection was limited to a small number of cells (4). This led us to speculate that our undifferentiated organoid-derived monolayers are heterogenous and that not all cells are susceptible to virus infection. Alternatively, the infection of permissive cells leads to the production of type I and/or III IFN, and IFN-induced proteins with antiviral activity inhibit the progression of the infection. To address the hypotheses, we first determined the cellular composition of organoid-derived monolayers using single-cell RNA-sequencing and found that the monolayer cells consist of a homogenous population of immature cholangiocytes, confirming our previous observations obtained using marker antibody staining. This finding demonstrates that RHDV can infect more cell types than previously thought, adding cholangiocytes to the list of RHDV-permissive cells along with primary hepatocytes (33). Cholangiocytes form bile ducts that channel bile from liver to the intestine (reviewed in (34)). If the virus also replicates in the intestine, it likely infects cells of the intestine before reaching the liver as it spreads via the faecal-oral route.

Next, we evaluated the IFN responses in organoid-derived monolayer cells using the small molecule IFN inhibitor Rux. This and similar inhibitors have previously been shown to increase virus replication in both continuous and organoid-derived cells (6, 10–12). To this end, we tested the pan-IFN inhibitor Rux and observed that it significantly improved virus replication as measured by both RT-qPCR and immunofluorescence staining, indicating productive viral genome replication and translation, respectively. Notably, significant effects were observed after adding Rux to cell culture media as late as 1 h after the infection. A similar effect was observed in intestinal organoid monolayers that were infected with human norovirus (6). Taken together, Rux boosts the replication of difficult to cultivate caliciviruses in organoid-derived cells.

In addition to improved virus replication, we observed an unexpected effect of Rux on our organoid-derived monolayers. While previous studies reported that Rux can induce apoptosis and suppress the proliferation of cancer cells (13, 35, 36), we observed a beneficial effect of Rux on our organoid-derived monolayers, with cells appearing to grow faster and attach more strongly to the surface of culture flasks. This stabilising effect is especially valuable for organoids and organoid-derived cells, as their growth characteristics can vary significantly, which affects the reproducibility of experiments (37). As we also observed variability in permissiveness between organoid cultures derived from different animals, we introduced a routine batch-testing in which cells derived from each rabbit were tested for RHDV susceptibility using immunofluorescence staining of infected cells and RT-qPCR (Supplementary Fig. 1).

To analyze the effect of Rux on IFN-induced gene expression in our organoid-derived monolayers, we used RT-qPCR to quantify the expression of *MX1*, *IFIT1*, and *GBP1*. In the absence of an infection, these genes were expressed at base levels while RHDV-infected cells expressed significantly higher levels. This indicates that RHDV triggers the IFN responses and may explain why the addition of Rux shortly after infection was sufficient to improve virus replication. However, the expression of all three genes was higher in cells infected in the absence of Rux compared to cells cultured with Rux, which confirms that Rux inhibits IFN responses in our system. The products of *MX1*, *IFIT1*, and *GBP1* are proteins with antiviral activity that are usually expressed along with many other IFN-induced proteins. More work is needed to determine whether the expression of any of these proteins affects RHDV replication.

Notably, *GBP1* expression is predominantly regulated by IFN-γ (type II IFN) (38), but expression can also be induced by type I IFNs ((39), reviewed in (40)). Since our monolayer cells cannot express IFN-γ, the observed expression of *GBP1* was likely due to type I IFN signalling.

Although virus growth in rabbit organoids was observed previously without Rux (4), attempts to passage RHDV were not successful. The ability of Rux to inhibit IFN-mediated signalling and stabilize organoid-derived monolayers improved the productivity of our organoids to a point that allowed us to passage lagoviruses repeatedly. It is likely that IFN inhibition is critical starting from passage two, where diluted supernatant that contains IFNs induced by the initial infection in passage one is used to infect the cells. The ability to productively grow and passage RHDV in cell culture will be useful for producing vaccines (to protect pet and farmed rabbits) and for selecting improved biocontrol agents. Previous studies that generated a pipeline for the selection of a new rabbit biocontrol agent were performed by passaging RHDV in laboratory rabbits, which is laborious and expensive (41).

This work demonstrates that RHDV replication in rabbit liver organoid-derived monolayers can be improved by supplementing the cell culture medium with the IFN inhibitor Rux. The inhibition of IFN-induced innate immune responses allows repeated passaging of RHDV. Moreover, Rux supports cell growth and improves monolayer cell attachment. Overall, this study improves our understanding of the developed liver organoid culture system and provides insights to increase its robustness.

## ACKNOWLEDGEMENTS

We thank the Australian Capital Territory (ACT) Parks and Conservation Services, the ACT Government, and contractors who conducted population control shooting of wild rabbits. We are grateful to Philip Hands (Black Mountain MicroImaging Centre, CSIRO) for providing advice and training in fluorescence microscopy.

## AUTHOR CONTRIBUTIONS

Conceptualization—T.S., M.F., I.S.; methodology— E.S., M.P., N.Y.G.P., O.F., M.J., T.S., M.F, I.S.; investigation—M.P., E.S., N.Y.G.P., O.F., I.S.; data curation—E.S., N.Y.G.P., O.F., M.J.; formal analysis—E.S., N.Y.G.P., T.S., I.S.; writing (original draft preparation)—E.S., M.F.; writing (review and editing)—E.S., M.P., N.Y.G.P., O.F., M.J., T.S., M.F., I.S.; visualization—E.S., M.J., M.F.; project administration—I.S.; validation—M.P., N.Y.G.P., O.F., E.S.; supervision—T.S., M.F., I.S.; funding acquisition—T.S.

## FUNDING

Funding for this work was provided by Meat and Livestock Australia (P.PSH.1059).

## ETHICAL STATEMENT

All animal samples used in this study were obtained opportunistically from wild rabbits that were shot for pest animal control purposes; animal ethics approval does not apply to this type of sampling in Australia.

## CONFLICT OF INTEREST

The authors declare no conflicts of interest.

## SUPPLEMENTARY MATERIAL

Here we provide an example of the organoid batch test procedure for rabbit liver organoid-derived monolayers. Each newly isolated batch of organoid monolayer cells were infected with RHDV1 and RHDV2, and the permissiveness of the cells was quantified using RT-qPCR and immunofluorescence staining.

**SUPPLEMENTARY FIG 1.**
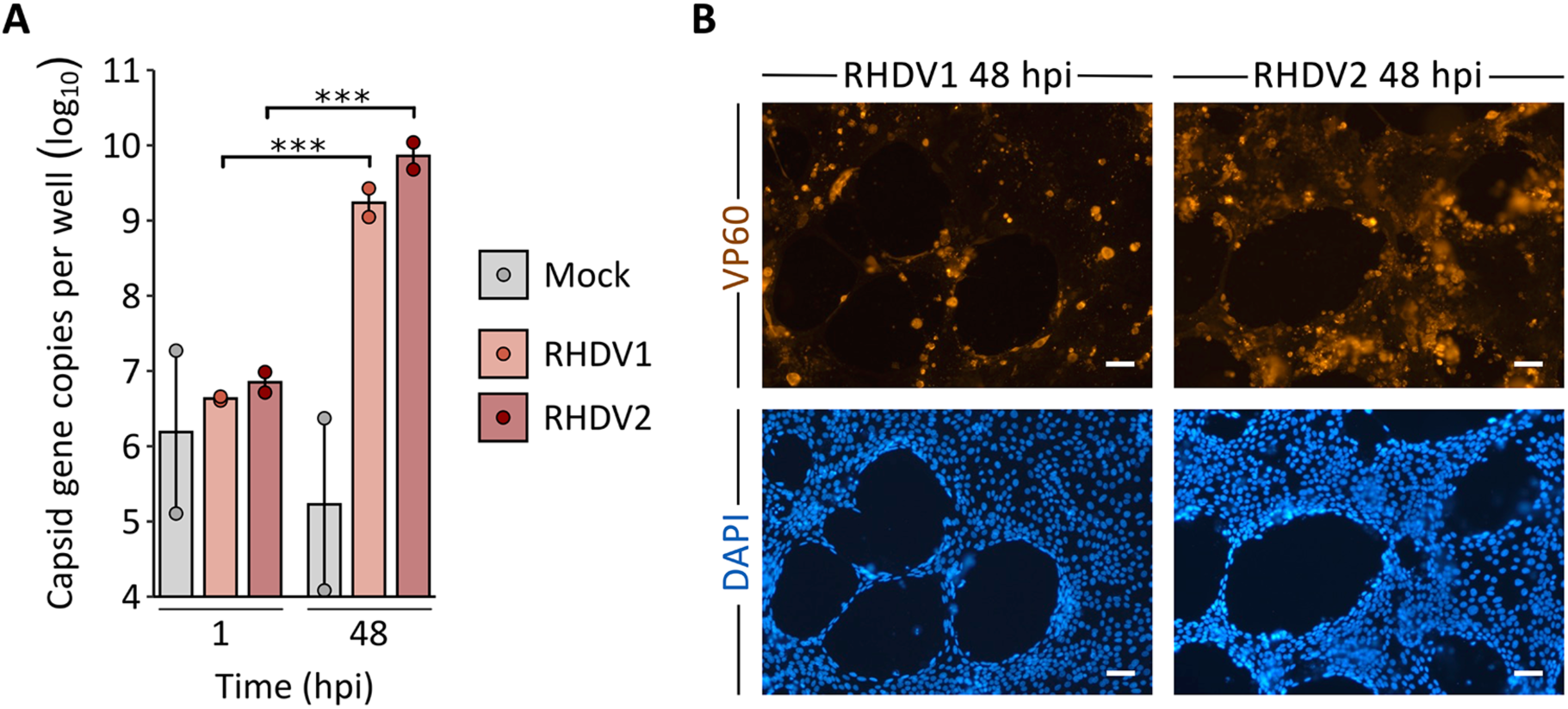
Infectivity test of liver organoid-derived monolayer cell batch designated WRL-6. (**A**) RT-qPCR was performed using RNA extracted from media after infection with RHDV1 and RHDV2; individual data points represent two biological replicates averaged from three technical replicates for each. Columns and error bars represent the mean values and the standard error of the mean, respectively. Significance is denoted by asterisks (\*\*\**p*<0.001) and was calculated using two-way ANOVA followed by Tukey’s HSD test. (**B**) The acetone-fixed cells were stained with antibodies to detect viral capsid protein VP60 (orange), DAPI was used to visualise cell nuclei (blue). Scale bar, 50 µM.

**SUPPLEMENTARY FIG 2.**
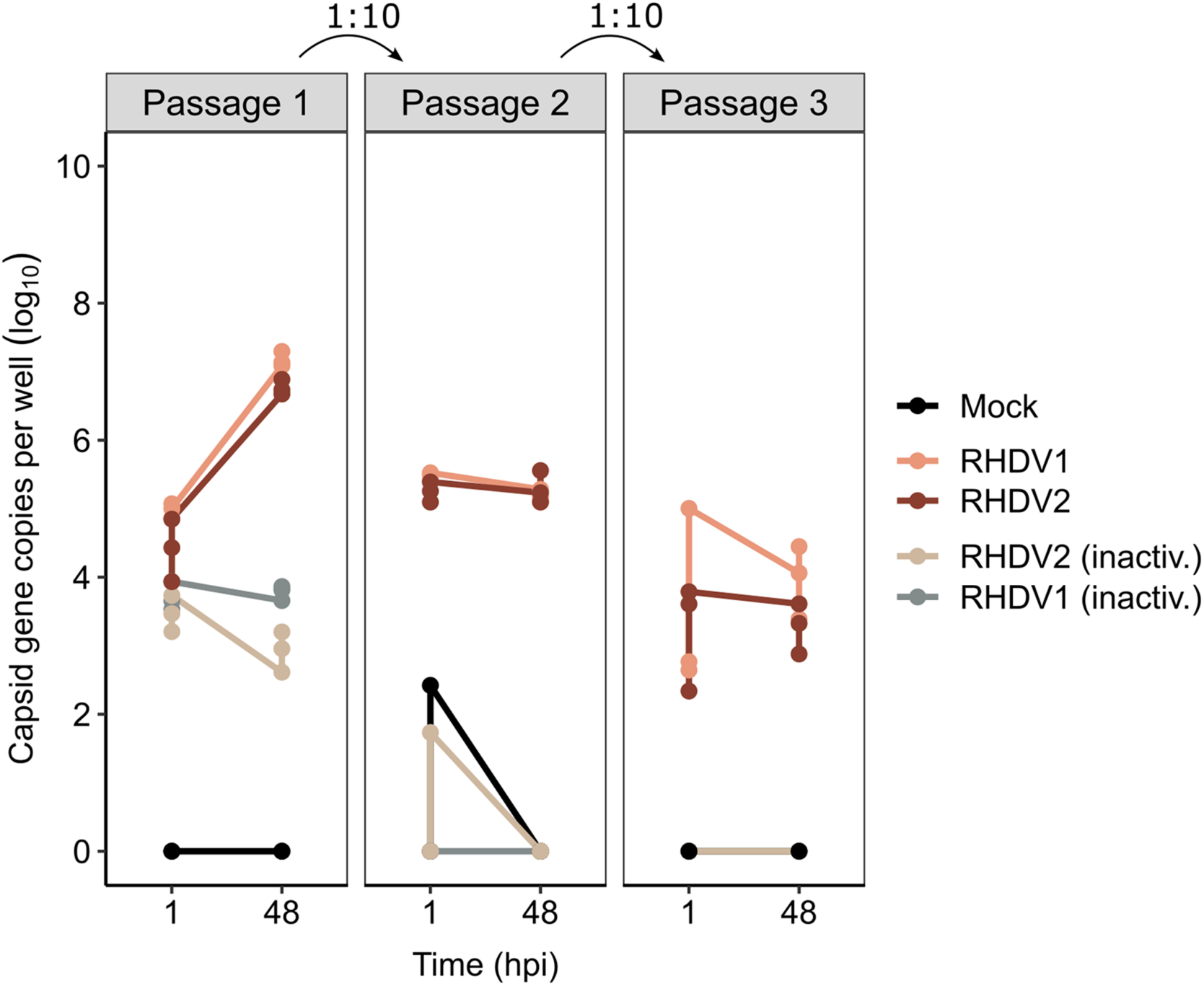
Passaging of RHDV in rabbit liver monolayers in the absence of Rux. The cells were infected with either RHDV1, RHDV2, heat-inactivated viruses (inactiv.) at a MOI of 0.01 or mock-infected, and the capsid gene copies were quantified at 1 or 48 hpi. The supernatant from passage one was diluted 1:10 and used as inoculum for passage two; passage three was infected in a similar manner. Individual data points represent three biological replicates averaged from two technical replicates for each.

**SUPPLEMENTARY TABLE 1.**
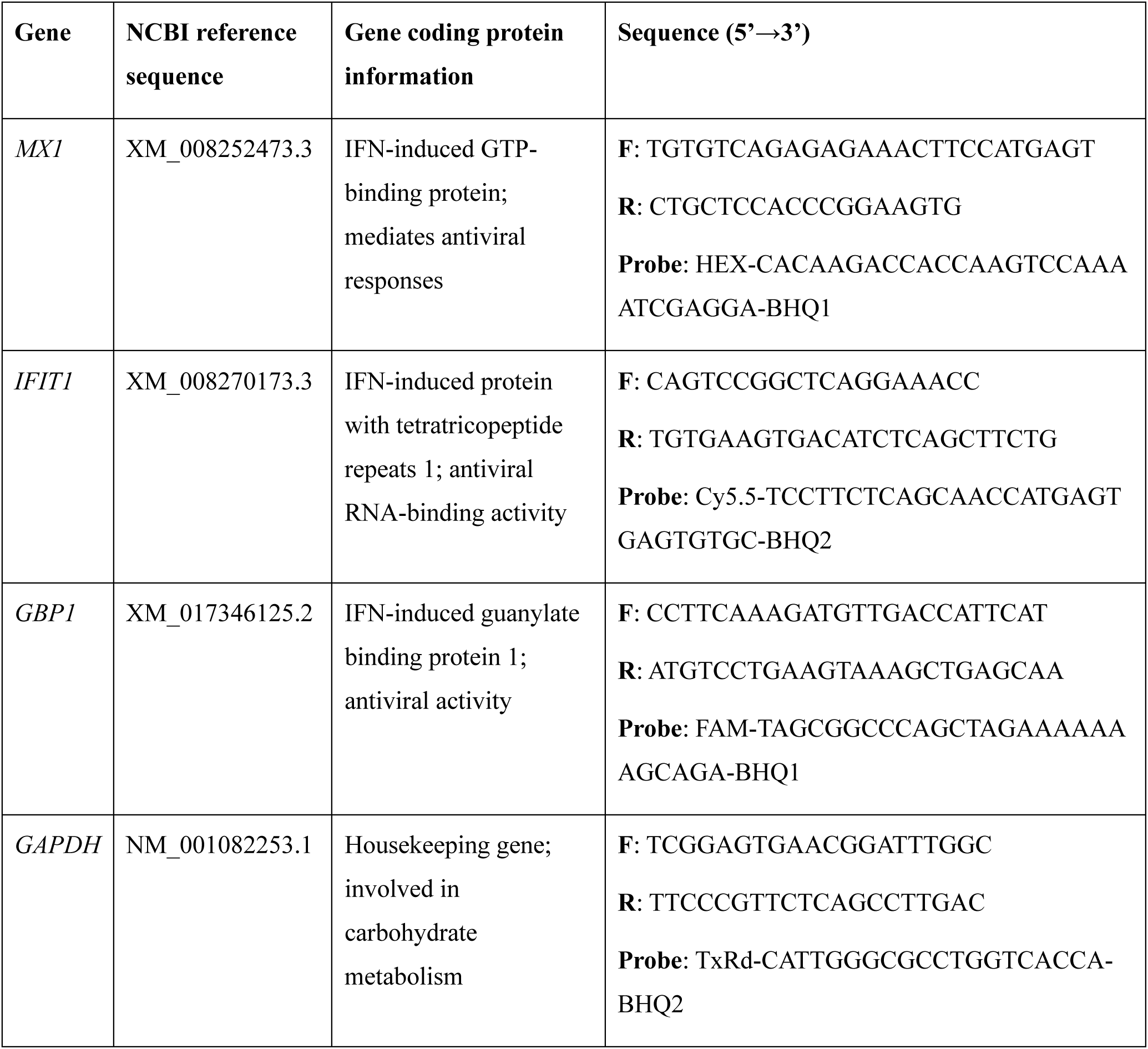
Primer sequences.

